# Migratory bats are sensitive to magnetic inclination during compass calibration

**DOI:** 10.1101/2023.05.03.539149

**Authors:** William T. Schneider, Richard A. Holland, Oskars Keišs, Oliver Lindecke

## Abstract

The Earth’s magnetic field is used as a navigational cue by many animals. For mammals, however, there is little data to show that navigation ability relies on sensing the natural magnetic field. In migratory bats, however, the calibration of a magnetic compass became plausible following experiments demonstrating a role for the solar azimuth at sunset in their orientation system. Here, we investigated how an altered magnetic field at sunset changes the nocturnal orientation of the bat *Pipistrellus pygmaeus*. We exposed bats to either the natural magnetic field, a horizontally shifted field (120°), or the same shifted field combined with a reversal of the natural value of inclination (70° to −70°). We later released the bats and found that the take-off orientation differed between all treatments. Bats that were exposed to the 120° shift were unimodally oriented northwards, in contrast to controls which exhibited a North-South distribution. Surprisingly, the orientation of bats exposed to both a 120°-shift and reverse inclination was indistinguishable from a uniform distribution. These results provide the missing link that these migratory bats calibrate a magnetic compass at sunset, and for the first time, they show that bats are sensitive to the angle of magnetic inclination.

## BACKGROUND

The question of how animals navigate as they migrate vast distances over varied terrain with ever-changing cue availability has interested scientists for centuries [1]. For migratory mammals the answers to this question are largely unknown. Long distance navigation appears even more remarkable for nocturnal migrants, which must find their way without the visual landmarks that are present during daytime [2]. This is the case for many migratory bats which travel hundreds of kilometres across Europe each year [3,4]. Whilst bats do possess the ability to echolocate, it is not suitable over distances greater than tens of metres [5], therefore it seems that further navigation tools must be necessary to successfully migrate. Night migrating birds are known to use the Earth’s magnetic field to aid their migratory navigation [6], but there has been no comprehensive evidence to show that migratory bats use the Earth’s magnetic field in the same way.

The soprano pipistrelle (*Pipistrellus pygmaeus*), is thought to migrate between north-east and south-west Europe [7]. In late summer they can be caught in great numbers on the Baltic coastline. Experimental releases of these bats have shown their take-off orientation can be measured to learn about their intended departure flight orientation [8]. Recently, it was found that adult soprano pipistrelles may calibrate compass information from the horizontal location, i.e., the azimuth of the setting sun [9], the first clear demonstration of such a mechanism to exist in animals. Earlier works in two non-migratory species of bats, *Eptesicus fuscus* and *Myotis myotis*, indicate that these animals relied on a magnetic compass to return to their home roosts at night [10,11]. However, it is not yet known for any species of migratory mammals whether a magnetic compass aids their long-distance navigation. Nevertheless, it is plausible that it is the Earth’s magnetic field which derives that compass calibrated in migratory pipistrelles. Therefore, in this study, we manipulated the magnetic field around bats during their sunset calibration to investigate whether this modified their take-off orientation when they were released later at night. If their nightly take-off orientation differed when shifts in the magnetic field were applied hours before, it would suggest that the Earth’s magnetic field is used in their sunset compass calibration.

## METHODS

### Experimental animals

Between the 20^th^ of August 2021 and the 10^th^ September, soprano pipistrelles (*Pipistrellus pygmaeus*) were caught at Pape Ornithological station, Latvia, using a large funnel trap [12]. Both males and females were caught as they appeared in the trap. The same night, the bats were checked for health and physical condition, aged, and then sorted by sex before being kept in darkness in wooden boxes for the calibration and release experiment the following night.

### Sunset exposure

On the night of the experimental release, kept bats were individually bagged to ensure darkness until the time of sunset exposure. The sex of bats was balanced for each treatment. Half an hour prior to sunset, bats were brought to the exposure sites and placed inside the sunset calibration cages [9]. These were put on a table 50 cm off the ground and oriented towards sunset. There were two exposure sites (figure 1*a*), one for control bats experiencing the natural magnetic field, and another for magnetically treated bats. The sites were approx. 100 m apart on dunes on the coast of the Baltic Sea, at Pape Ornithological station. The sites were not in view of each other due to the vegetation. Half an hour after sunset bats were returned to bags and carried to a room to be given ID’s (to which the experimenter was blind), and then individually bagged again.

**Figure 1.**
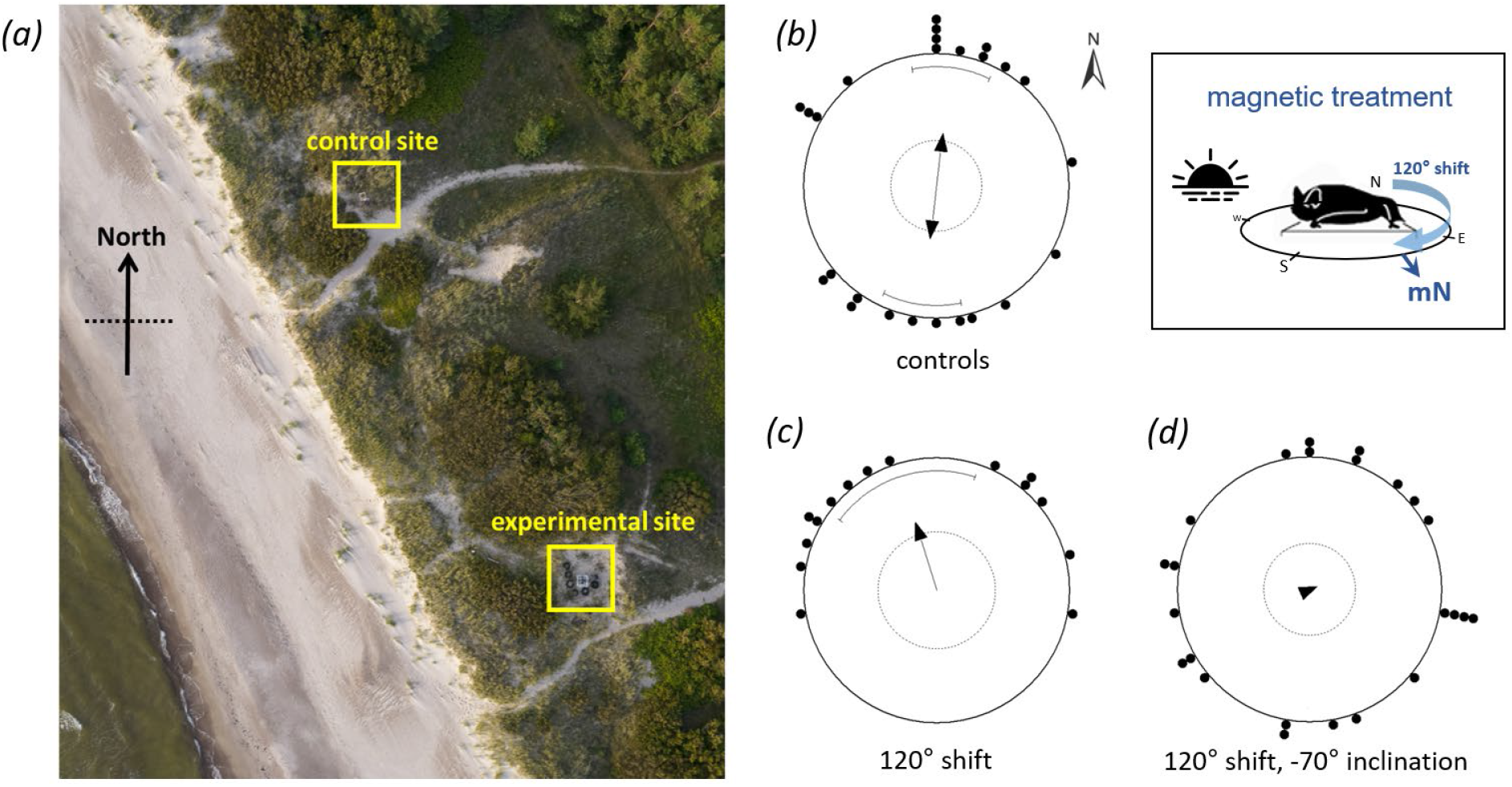
Sunset calibration sites (a) for control bats which experienced the natural magnetic field, and for experimental bats which experienced manipulated magnetic field conditions within a Helmholtz coil. Take-off orientations for control bats (b), 120° horizontally shifted bats (c), and bats who experienced the horizontal 120° shift combined with a reversed magnetic inclination of −70° (d).

### Yurt release

After the blinding procedure, individually bagged bats were placed inside a thermally lined box with a hot water bottle to prevent torpor. Releases were conducted inside a Mongolian yurt, also on the Pape Ornithological station site. The yurt was circular, 5.6m in diameter, with a height of 2.4 m at the centre and 1.5 m at the edge. In the centre of the yurt a circular release box, CRBox [9], was used to conduct the releases. The experimenter outside the yurt released bat individually using a string pulley system. Upon release, a narrowband ultrasound detector (Pettersson D-100) was used to listen for any echolocation that signified take-off. Bats were given 3 minutes to take-off. Once this time had elapsed, or when a take-off had been audibly detected, the experimenter entered the yurt, the bat was re-caught, and the orientation of its take-off was recorded. Bats were then released into the wild.

### Magnetic treatment

Control animals experienced the natural magnetic field of Pape Ornithological station (51.6 latitude, 21 longitude). The magnetic field parameters for August 2021 given by the International Geomagnetic Reference Field (IGRF) 13^th^ generation for this location are: H = 16728, Z = 48432, F = 51239, I = 70.9, and D = 7.3. Manual measurements of the natural magnetic total intensity and inclination were made before every experiment using a 3 Axis Milligaus Meter (Model MR3, Alpha Lab, Inc.) magnetometer, the mean and standard deviation of these were: F = 51126 nT ± 71, I = 71.17° ± 0.48. Manipulations to the magnetic field for the magnetically treated bats were made using a single-wrapped, three-axis Helmholtz coil (Claricent, Munich; 1% homogeneity in a 60 cm diameter). One magnetic treatment was conducted at a time, with control bats being tested simultaneously on the same night. Once completed, the next magnetic manipulation experiment began, with continued testing of control bats. For the inclination experiment, bats were placed within a magnetic field shifted clockwise horizontally 120° during sunset calibration. All other magnetic parameters were kept constant. Across 8 experimental nights, the mean and standard deviation of magnetic total intensity and inclination inside the coil were: 51105 nT ± 88 and 71.05° ± 0.37. For the shift and inclination reverse experiment, bats were again placed within a magnetic field shifted clockwise horizontally 120°, and this time the Z component of the field was reversed, producing a negative inclination. This experiment lasted 4 nights, across which the mean and standard deviation of magnetic total intensity and inclination inside the coil were: 51138 nT ± 125 and −70.9° ± 0.08. Magnetic locations on Earth that share the conditions created by both experiments can be found in the electronic supplement (ESM1).

### Statistical methods

Mean bearings and vector lengths were calculated using Oriana 4.02 (Kovach Computing Services). The Rayleigh test was used to test for unimodal non-uniformity of circular distributions. The test suggested a non-unimodal distribution in both the control and the inclination group. To specifically describe the patterns of orientation in these groups, we followed a likelihood-based modelling approach (package CircMLE, R version 3.5.2) which allows comparison of circular data with multiple potential models of orientation [13,14]. For each experimental group, resulting models were then compared by means of the corrected Akaike information criterion (AICc) and the corresponding model weights (see the electronic supplementary material for details, ESM2). The Mardia-Watson-Wheeler test was applied to test for significant differences between group orientations. To compare the directedness based on the r-values between the groups, we applied a bootstrap technique that enables a comparison of non-unimodally oriented groups with a significantly oriented group [15] (see ESM2).

## RESULTS

The orientation of control bats, who experienced the natural magnetic field during sunset calibration, was homogeneously axially bimodal Northwards and Southwards (figure 1*b*), p = 0.016 (Rayleigh’s test: n = 26, Z = 4.126, mean vector µ_axial_ = 4.5°, r = 0.398; see the electronic supplementary material, ESM2, for details of the likelihood-based modelling approach). Bats which experienced a field shifted 120° clockwise during sunset calibration had take-off orientations towards NNW (µ = 342.3°), p = 0.01 (n = 15, Z = 4.382, r = 0.54). Finally, bats which were given a 120° shifted field as well as a reversed inclination (−70.9°), did not show any clear orientation at take-off, and were indistinguishable from a uniform distribution (p = 0.917, n = 24, Z = 0.089, µ = 66.9°, r = 0.061; see also ESM2). This group was significantly different from the orientation of the group that was exposed to the horizontally shifted field (+70.9° inclination), p = 0.041 (MWW test, W = 6.39, df = 2).

## DISCUSSION

We found that two different manipulations of the Earth’s magnetic field during sunset differentially effect the take-off orientation of migratory bats released later at night. Because the control bats flew bimodally North and South, it is difficult to assess if the changed orientation of the shifted bats corresponds directly to the degree of the horizontal shift of the magnetic field. However, the unimodal Northwards orientation of the shifted bats represents a dramatic shift in behaviour from the control bats suggesting that the bats are able to sense the magnetic field shift. Furthermore, the random distribution of the inclination-reversed bats is a contrasting result to both the control and shifted bats. Significantly, these results provide the first evidence that the Earth’s magnetic field is used for orientation by migratory bats, and specifically we have shown that sunset calibration is likely to be utilising a magnetic compass.

The bimodality of the control bats has not been observed in previous studies of *P*.*pygmaeus*, but has in *P. nathusii* at the same location [8]. In late summer, the generally expected migratory direction of *P*.*pygmaeus* is south towards central Europe [8], however there is still little known about the migratory route of this bat species and it is possible that two migratory routes are used; one following the coast southwards, and another that goes northwards into Scandinavia and only then proceeds in a south westerly direction [16]. It is also possible that there is a mix of resident and migratory individuals at the field site, although if that were the case then we might not expect such a clear unimodal preference for North in the shifted bats. In migratory birds it is frequently observed that a proportion of animals will exhibit a reversed migratory orientation [17,18]. Interestingly, the behaviour of the shifted bats suggests that the magnetic treatment had a unifying effect, reverting the preference of all bats to a common direction, in this case North; the opposite of the generally expected migratory direction.

It has been observed in a species of migratory bird that inclination reversal (without a horizontal shift) reversed their orientation [19,20]. If the effect of any magnetic manipulation were to reverse migratory direction, perhaps due to stress or confusion, then we would expect the same to be the case for the shifted-and-inclination-reversed treatment group. However, we found that the orientation of the inclination-reversed bats appeared to be random. This difference between the shift and the shift with reversed inclination is suggestive of the bats being able to detect differences in the horizontal and vertical components of the magnetic field, and therefore an ability to sense magnetic inclination. The combination of magnetic total intensity and inclination that the inclination-reversed bats experienced can naturally be found in the Southern Indian Ocean (see ESM2). If the altered declination resulting from the shift is also considered, then there are no possible locations where these magnetic parameters occur naturally. Therefore, whilst the shifted bats experienced only a rotated field, the shifted-and-inclination-reversed bats experienced a magnetic field very different from anything they would normally experience which may explain the lack of any directionality.

Significantly, the magnetic manipulations that we applied occurred only during the sunset period. Bats were later released in a natural magnetic field, with all other environmental cues obscured. The effects of the magnetic manipulations are therefore long-lasting, modifying bat behaviour hours after they were removed from the altered magnetic field conditions. This suggests either that the sunset period is key to calibrating a magnetic compass, or in the case that a magnetic compass is not being utilised, that regardless, magnetic fields are responsible for producing a long lasting behavioural reaction in migratory bats.

## ETHICS STATEMENT

All work was conducted under the permit no. 21/2021 to the Institute of Biology, University of Latvia. All experimental procedures were conducted in accordance with national and local guidelines for the use of animals in research.

## DATA ACCESSIBILITY

The data are provided in the electronic supplementary material.

## ACKNOWLEDGEMENTS

We thank Ivo Dinsbergs for the drone picture of the experimental sites and Donāts Spalis (both University of Latvia) for supporting our work at PBRS. We are grateful to the voluntary station crews who helped with bat catching and animal care, particulary to Gunārs Pētersons, Viesturs Vintulis, Valts Jaunzemis and Roberts Jansons.

## REFERENCES

1. Darwin C. 1873 Perception in the Lower Animals. Nature 1873 7:176 7, 360–360. (doi:10.1038/007360c0)

2. Martin GR. 1990 The Visual Problems of Nocturnal Migration. Bird Migration, 185–197. (doi:10.1007/978-3-642-74542-3_13)

3. Pētersons G. 2004 Seasonal migrations of north-eastern populations of Nathusius’ bat Pipistrellus nathusii (Chiroptera). MYOTIS 41–42, 29–56.

4. Hutterer R, Ivanova T, Meyer-Cords C, Rodrigues L. 2005 Bat Migrations in Europe A Review of Banding Data and Literature. Bonn: Federal Agency for Nature Conservation.

5. Stilz W-P, Schnitzler H-U. 2012 Estimation of the acoustic range of bat echolocation for extended targets. The Journal of the Acoustical Society of America 132, 1765. (doi:10.1121/1.4733537)

6. Bingman VP. 1987 Earth’s Magnetism and the Nocturnal Orientation of Migratory European Robins. The Auk 104, 523–525. (doi:10.2307/4087555)

7. Sztencel-Jabłonka A, Bogdanowicz W. 2012 Population genetics study of common (Pipistrellus pipistrellus) and soprano (Pipistrellus pygmaeus) pipistrelle bats from central Europe suggests interspecific hybridization. Canadian Journal of Zoology 90, 1251–1260. (doi:10.1139/Z2012-092/ASSET/IMAGES/Z2012-092TAB4.GIF)

8. Lindecke O, Elksne A, Holland RA, Pētersons G, Voigt CC. 2019 Orientation and flight behaviour identify the Soprano pipistrelle as a migratory bat species at the Baltic Sea coast. Journal of Zoology 308, 56–65. (doi:10.1111/JZO.12654)

9. Lindecke O, Elksne A, Holland RA, Pētersons G, Voigt CC. 2019 Experienced Migratory Bats Integrate the Sun’s Position at Dusk for Navigation at Night. 29, 1369–1373.e3. (doi:10.1016/J.CUB.2019.03.002)

10. Holland RA, Borissov I, Siemers BM. 2010 A nocturnal mammal, the greater mouse-eared bat, calibrates a magnetic compass by the sun. Proceedings of the National Academy of Sciences of the United States of America 107, 6941–6945. (doi:10.1073/PNAS.0912477107/ASSET/65092644-5B68-451A-BA7F-069E2FAA7ED2/ASSETS/GRAPHIC/PNAS.0912477107FIG03.JPEG)

11. Holland RA, Thorup K, Vonhof MJ, Cochran WW, Wikelski M. 2006 Bat orientation using Earth’s magnetic field. Nature 2006 444:7120 444, 702–702. (doi:10.1038/444702a)

12. Keišs O, Spalis D, Pētersons G. 2021 Funnel trap as a method for capture migrating bats in Pape, Latvia. Environmental and Experimental Biology 19, 7–10-7–10. (doi:10.22364/EEB.19.02)

13. Fitak RR, Johnsen S. 2017 Bringing the analysis of animal orientation data full circle: model-based approaches with maximum likelihood. Journal of Experimental Biology 220, 3878–3882. (doi:10.1242/jeb.167056)

14. Schnute JT, Groot K. 1992 Statistical analysis of animal orientation data. Animal Behaviour 43, 15–33. (doi:10.1016/S0003-3472(05)80068-5)

15. Chernetsov N, Pakhomov A, Kobylkov D, Kishkinev D, Holland RA, Mouritsen H. 2017 Migratory Eurasian Reed Warblers Can Use Magnetic Declination to Solve the Longitude Problem. Current Biology 27, 2647–2651.e2.

16. In press. Identifying migratory pathways of Nathusius’ pipistrelles (Pipistrellus nathusii) using stable hydrogen and strontium isotopes - Kruszynski - 2021 - Rapid Communications in Mass Spectrometry - Wiley Online Library. See https://analyticalsciencejournals.onlinelibrary.wiley.com/doi/10.1002/rcm.9031 (accessed on 10 March 2023).

17. Akesson S. 1999 Do passerine migrants captured at an inland site perform temporary reverse migration in autumn? Ardea,

18. Sandberg R, Moore FR. 1996 Migratory orientation of red-eyed vireos, Vireo olivaceus, in relation to energetic condition and ecological context. Behavioral Ecology and Sociobiology 1996 39:1 39, 1–10. (doi:10.1007/S002650050261)

19. Wiltschko W, Munro U, Ford H, Wiltschko R. 1993 Magnetic inclination compass: A basis for the migratory orientation of birds in the Northern and Southern Hemisphere. Experientia 1993 49:2 49, 167–170. (doi:10.1007/BF01989423)

20. Wiltschko W, Wiltschko R. 1972 Magnetic Compass of European Robins. Science 176, 62–64. (doi:10.1126/science.176.4030.62)

